# Can urban fence lizards identify cats?

**DOI:** 10.64898/2026.07.22.735083

**Authors:** Emma M. Pfister, Daniel T. Blumstein, H. Bradley Shaffer, David R. Daversa

## Abstract

Urban habitats can serve as wildlife refuges, or as ecological traps. As global urbanization continues to increase, native prey species may experience reduced threat from natural predators that tend to avoid human-populated areas. However, such protection often comes at the expense of susceptibility to synanthropic predators, one of the most prevalent being domestic house cats (*Felis catus*). While cats are a known driving force of mortalities in urban wildlife, their sublethal effects remain understudied. To better understand the behavioral effects of synanthropic predators on native urban prey, we examined responses of western fence lizards (*Sceloporus occidentalis*) to Artificial Intelligence (AI)-generated visual stimuli of native predators (snakes) and synanthropic predators (house cats). Visual stimuli included: California striped racers (*Masticophis lateralis*), a native predator; domestic cats, a synanthropic mammalian predator (*Felis catus)*; and desert cottontails (*Sylvilagus audubonii*), a native mammalian non-predator. We quantified lizard responses as the change in rates of looking and locomotion during and following the video presentation. Our results revealed that lizards discriminated cats from the control by looking more within the first 90 seconds following exposure. Lizards looked more across the entire 5-minute observation period following exposure to all stimulus types compared to control conditions, indicating that AI generated visual stimuli effectively elicit behavioral responses. Together, these findings indicate that urban lizards initially detect and respond to visual cues of cats, suggesting that cat-induced mortality is not a result of inability to detect such predators. Thus, cats influence urban prey not only through direct predation risk, but also changes in behavior associated with the presence of synanthropic predators.

## Introduction

As urbanization alters natural landscapes, it is becoming increasingly important to understand the effects of human development on ecological systems. Habitat loss and fragmentation often lead to reduced native predation pressure in urban environments, altering predator-prey dynamics (Seto et al. 2012; Grimm et al. 2008; Moller et al. 2012; Muhly et al. 2011). The decline of natural predators in increasingly urban habitats may provide increased protection to organisms that are typically preyed upon in rural areas (‘the urban refuge hypothesis’, Moller et al. 2012; Rebolo-Ifrán et al. 2017). While such environments can serve as refuges, they may also act as ‘ecological traps’ if native prey populations are forced to coexist with synanthropic predators that thrive in human-modified habitats (Tomialojc 1970; Johnston et al. 2001; Moller et al. 2012; Salo et al. 2007; Loss et al. 2013).

Domestic house cats (*Felis catus*) are a synanthropic species that poses a significant predation risk to a broad range of wildlife in urban areas (Loss et al. 2013). Cat predation in the United States accounts for an estimated 6.3 to 23.3 billion small bird and mammal mortality rates annually (Loss et al. 2013). The diet of free-roaming domestic cats is influenced by locally available prey, resulting in the consumption of substantial amounts of small wildlife (Lepczyk et. al. 2023). Domestic cats were first brought to North America around 450 years ago (Welker et al. 2025), and their effects on urban biodiversity are enormous (Magle & Crowther 2023). Despite their recognition as a notable mortality risk to urban prey, there is little known about the sublethal effects of the species, including impacts on behavior. Changes in wildlife behavior can be one useful assay for the ecological significance of a stressor, including novel predation (Greggor et al. 2019).

Animals detect predators using a variety of modalities, including visual cues (Caro 2005). Antipredator vigilance is often used in visual discrimination experiments to quantify risk assessment and changes in looking behavior after being exposed to a threatening stimulus (e.g., Blumstein et al. 2000; Kong et al. 2022). It is important to control for a variety of features when studying stimulus discrimination. Using live animals creates novel welfare issues (for both the experimental subjects and the models), an issue that can be avoided by using video playback. While video playback has been used to study opponent assessment (Ord & Evans 2002), there is a long history of taxidermic mounts used for studies on predator recognition (Blumstein et al. 2000; Carlson et al. 2017, Tinbergen 1989; Curio et al. 1978). However, lizards exposed to taxidermic mounted cats do not engage in natural behavior (Lishmund et al. 2025). An additional shortcoming of taxidermic models is that often only one or a few mounts are used in an experiment, which then limits the inference to the mount rather than to the predator type. Until now, studies using visual predator cues have been limited by the inability to create controlled replicates for each stimulus. However, recent advances in artificial intelligence (AI) now permit the creation of a collection of replicated visual stimuli to simulate ecologically relevant, yet controlled, encounters. This approach allows one to standardize the background and movement while varying the stimulus in a controlled way. Thus, differential responses can be attributed to the stimulus characteristics alone rather than confounding variables associated with specific videos.

To examine the behavioral effects of cats on urban prey, we used AI-generated videos to expose western fence lizards (*Sceloporus occidentalis*) to visual stimuli of native predators, synanthropic cat predators, and native non-predators. Stimulus types included California striped racers (Colubridae: *Masticophis lateralis*) as native predators, domestic cats (Felidae: *Felis catus*) as synanthropic predators, and desert cottontails (Leporidae: *Sylvilagus audubonii*) as native mammalian non-predators. Because western fence lizards monitor visual stimuli and co-exist with domestic cats in the area where we studied them (Southern California, USA), we hypothesized that subjects would perceive cats as a threat and alter their behavior in comparison to control and non-predatory stimulus conditions.

## Materials and Methods

### Study Species and Site

Western fence lizards are widely distributed across Western North America, making them an abundant and easily accessible species for study (Bishop et al. 2023). As ectotherms, they occupy a broad range of habitats with high levels of sun exposure, such as grasslands, woodlands, and chaparral, and have been effectively used as model organisms in a wide range of studies on environmental change and urbanization (Bishop et al. 2023; Buckley et al. 2008; Sinervo et al. 2010; Putman et al. 2019; Putman et al. 2025; Putman et al. 2024). They also live in and around cities, including Los Angeles, CA (USA), where they are exposed to cats, making them a suitable focal species for our study.

This study took place on the University of California Los Angeles (UCLA) campus under approved IACUC protocols (ID: ARC-2025-016) and state collection permits (SC-2480). We collected sixteen wild adult western fence lizards from a single population in Sage Hill, a natural reserve on the UCLA campus. The lizards had no previous history of rearing or use in experiments. Upon capture, we measured snout-vent length (mm) and weight (g). We transported lizards in cloth bags to an experimental area located in the UCLA Mathias Botanical Garden. Subjects were housed individually in four separate outdoor enclosures (1.52 m x 0.61 m x 0.91 m) constructed from wood and metal hardware cloth. We installed a clear plexiglass panel (47.5 cm x 47.5 cm) onto the short side of each enclosure to allow monitors to be placed flush against the enclosure and ensure visibility (Figure S1). We fastened opaque cloth barriers to all enclosures to prevent visual interactions between individuals and to prevent monitor glare (Figure S1). Enclosures contained a sand substrate covering the base, two wooden hiding structures, and a brick for basking (Figure S1). We visually inspected lizards daily for signs of injury, illness, or death and fed them one cricket per day. We housed lizards in enclosures for an overnight acclimation period of 24 hours prior to trials.

We conducted trials over a period of four weeks, with four new subjects tested per week (n = 16). We used a within-subjects, Latin square design, where each lizard experienced all four stimulus types, one per day (Table S1). This design ensured that each lizard was exposed to all stimuli while minimizing potential order effects.

### Stimulus Videos

We created four, 15 s long videos for each stimulus type (snake, cat, and rabbit) using Google Veo 2 (Google, 2025) to ensure biologically accurate movement and features of each species. The four unique videos per stimulus type depicted slightly different individuals of the same species displaying similar movement patterns. The animals were scaled to fill the frame to ensure that all behavioral responses could be attributed to species-specific visual characteristics rather than differences in size. All stimuli were superimposed onto a standardized background image of Sage Hill (Figure S2) and were introduced through faded transitions to avoid overly rapid visual changes. We used a static image of the Sage Hill background with no animal present as the control. All stimulus videos are contained in Supplementary Material (Videos S1 - S12).

We presented visual stimuli on an 18.5-inch portable IPS monitor (CoolHood; 1920 x 1080 (FHD) resolution, 120 Hz refresh rate, 100% sRGB color gamut). Although this display lacked ultraviolet (UV) wavelengths detected by lizards (Fleishman et al. 2024), it served as an optimal alternative to cathode ray tubes (CRT) used in past studies on exposure to visual stimuli (Macedonia et al. 1993). There is evidence that visual stimuli can appear distorted to subjects when presented on CRT screens which tend to have lower refresh rates compared to LCD technology (Railton et al. 2010). Because lizards have high critical flicker fusion frequencies (Jenssen & Swenson 1974), using an IPS monitor with a high refresh rate ensured that visual stimuli would be clearly presented to subjects.

### Experimental Design

Prior to the start of each trial, we placed monitors against the plexiglass panel of the enclosure and mounted a camera on the edge of the enclosure. We lifted the roof of the enclosure to make sure that we could visibly observe the subject. After a five-minute acclimation period following experimental setup, we displayed the static image of Sage Hill with no stimulus present for five minutes (Figure 1). We then presented the 15 s stimulus video (snake, cat, rabbit, or control) which transitioned into the static image of Sage Hill for an additional five minutes before the monitor was turned off. Observations continued for five minutes after the monitor was turned off (Figure 1). The observer (E. Pfister) quietly dictated behavioral transitions into a digital recorder starting 1 min prior to the stimulus onset and for 10 min following stimulus exposure. The total focal observation time per individual was 11 minutes and 15 seconds, consisting of 1 minute pre-stimulus, 15 seconds stimulus, and 10 minutes post-stimulus. Our ethogram (Table S2) contained vigilance, locomotion, and display behaviors. We classified all behaviors as either horizontal (on the substrate) or vertical (on the sides of the enclosure). Display behaviors consisted of push-ups, head bobs, and tongue flicks. We scored individuals as “out of sight” at any time in which they were no longer visible.

**Figure 1.**
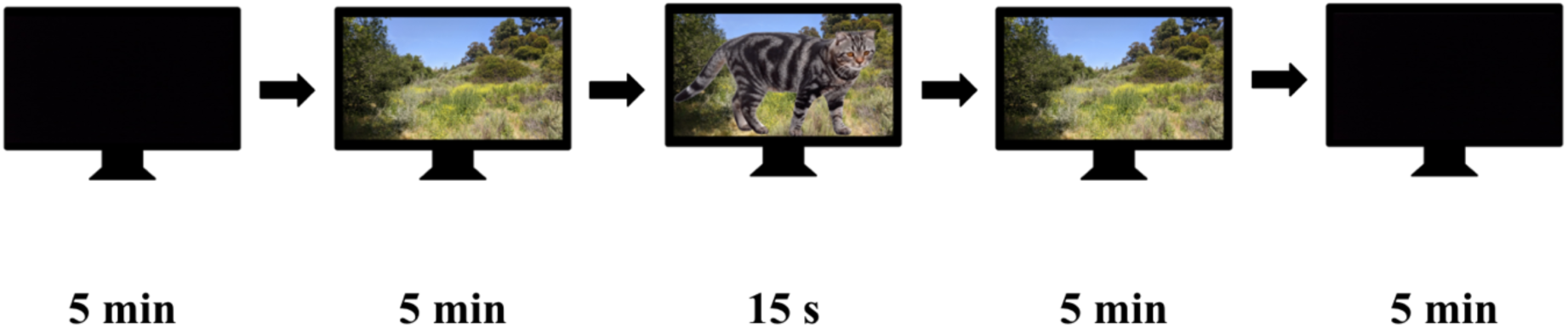
Visual stimulus sequence. All stimuli – cats, snakes, rabbits - were superimposed onto a standardized background image of Sage Hill and were introduced through faded transitions We presented visual stimuli on an 18.5-inch portable IPS monitor for 15 seconds, preceded and followed by a period of 5 minutes with no image and 5 minutes with the Sage Hill background.

### Statistical Methods

We quantified focal observations using the behavioral event recorder JWatcher 1.0 (Blumstein & Daniel 2006). All behaviors recorded in JWatcher were defined as mutually exclusive. We separated focal observations per trial into a 60 s pre-stimulus baseline, followed by ten 30 s time intervals. Focal observations for the 15 s stimulus exposure were included in the first 30 s time interval.

We analyzed focal observations in R (R Core Team 2025) using RStudio (Posit Team 2026) where we calculated behavioral rates per time interval as the quantity of behaviors divided by the total time in sight within each time interval. To determine the changes in behavioral rates over time, we subtracted 60 s pre-stimulus baseline rates from the behavioral rate from each time interval.

To examine the change in rate of key behaviors over time following stimulus exposure, we fitted general linear mixed models (GLMMs) with a Gaussian error structure to model the change in rates of horizontal looking, vertical looking, horizontal locomotion, vertical locomotion, combined looking, and combined locomotion. We fitted our initial model for the first 60 s and the first 90 s using the lme4 (Bates et al. 2015) and lmerTest (Kuznetsova et al. 2017) packages in R, which analyzed immediate responses to visual stimuli. To examine broad behavioral responses over time we also fitted our model for the entire post-stimulus time period of 5 minutes. We fitted all initial models with an interaction of treatment and time and included lizard ID as a random effect to account for repeated measures. We observed no significant interaction between treatment and time for any behavior in any given model (all p > 0.05), indicating no differential treatment effect as time went on following stimulus exposure. Therefore, we simplified all models to include main effects only.

We also explored the effects of other potentially influential factors, including: trial number (representing trial order) to account for possible habituation, body length (mm) or body mass (g) to account for possible size-related effects on behavioral responses (Preiser & Orrock 2012; Baxter-Gilbert et al. 2018), sex, time of day, and cage. our sample size did not permit a single model including all the above factors, and so we fitted different models for each factor. We evaluated body length and weight in separate models due to strong correlation between the two variables. We observed that body length had a significant effect in the model of vertical locomotion and retained the factor as a covariate in the corresponding final models (Figure S3). All other factors were not influential and therefore were excluded from all final models.

Our final models were fitted for changes in rates for horizontal looking, vertical looking, horizontal locomotion, and vertical locomotion. We examined these rates across two time intervals: 90 seconds and 5 minutes. We did not analyze combined looking and locomotion in final models because we found that aggregation of behaviors decreased interpretability. We did not find significant behavioral changes within the first 60 seconds following stimulus exposure and therefore focused our analysis on the first 90 seconds. We calculated post-hoc pairwise comparisons of specific treatments with Tukey HSD using the emmeans package in R (Lenth & Piaskowski 2026) and plotted the marginal mean treatment effect. We checked assumptions (Figure S4 - S11) for all final models using the performance package in R (Lüdecke et al. 2021).

## Results

Lizards modified the rate at which they looked (horizontally) within the first 90 s following stimulus exposure. This change in rate was influenced by stimulus type (ANOVA: F = 3.86, p = 0.011; Figure 2A). Specifically, cats elicited increased rates of horizontal looking relative to control conditions (t = −3.28, p = 0.007, Figure 2A), but other stimuli did not (all p > 0.05; Figure 2A).

**Figure 2.**
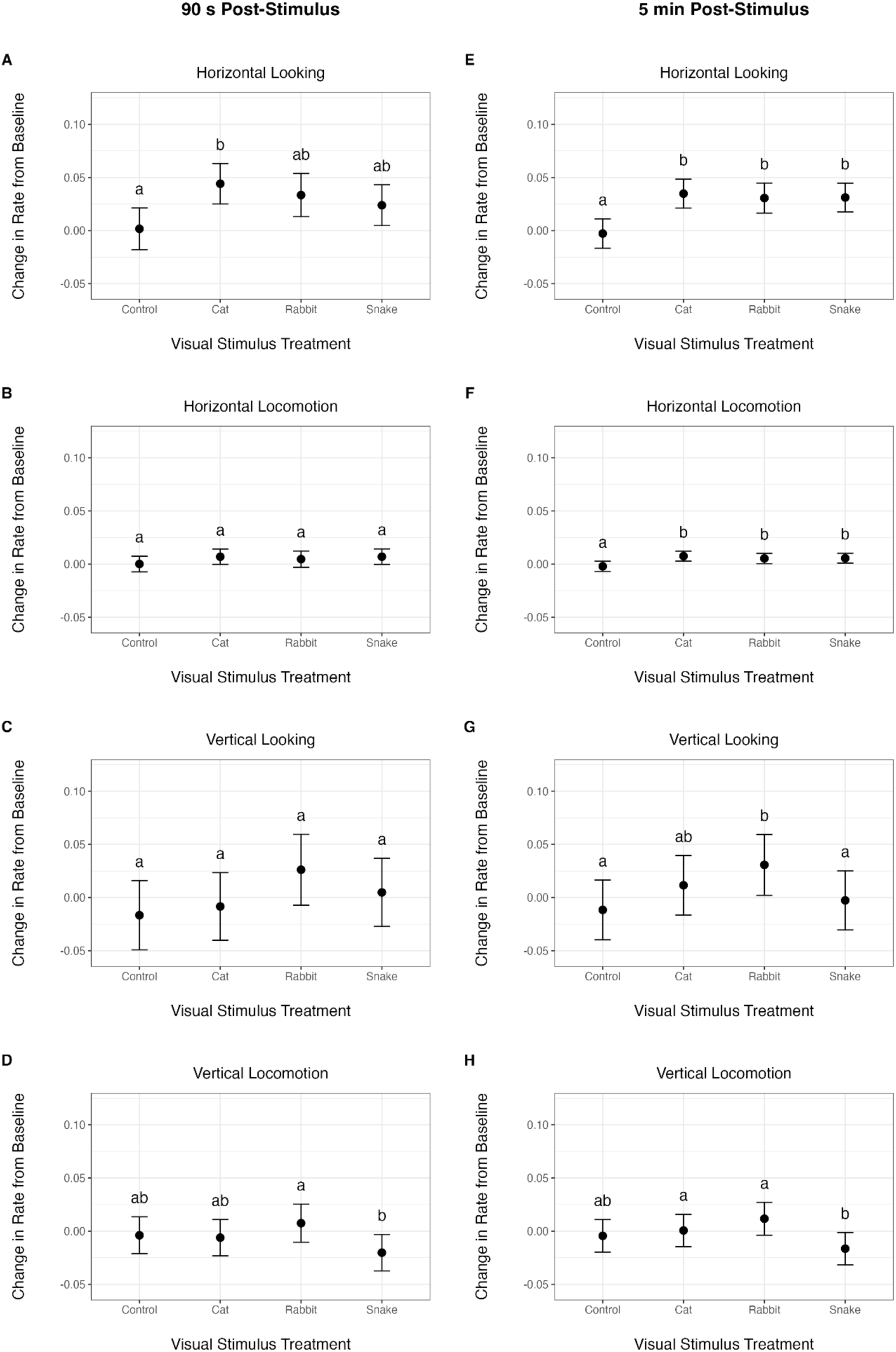
The vigilance and locomotory responses of western fence lizards to experimental exposures of cats, rabbits, and snakes. Panels depict estimated marginal means for changes in rates of horizontal looking (A, E), horizontal locomotion (B, F), vertical looking (C, G), and vertical locomotion (D, H) for the first 90 s (A-D) and 5 min (E-H) post-stimulus. Points represent mean changes in rates per stimulus type (control, cat, rabbit, and snake) relative to baseline behaviors. Error bars represent ±95% confidence intervals. Letters represent statistically significant differences between stimulus treatment groups evaluated using a Tukey contrast (see text).

Lizards modified their vertical locomotion behavior within the first 90 seconds following stimulus exposure as a function of stimulus type (ANOVA: *F* = 2.67, *p* = 0.049; Figure 2D). Specifically, rabbits elicited higher rates of vertical locomotion than snakes (*t* = 2.81, *p* = 0.029; Figure 2D). No stimulus type elicited significant differences in vertical locomotion relative to control conditions (*p* > 0.05 in all cases; Figure 2D). Stimuli did not influence changes in horizontal locomotion (ANOVA: *F* = 1.03, *p* = 0.381; Figure 2B) or vertical looking rates (ANOVA: *F* = 2.10, *p* = 0.103; Figure 2C) within the first 90 seconds following stimulus exposure.

Across the entire 5-min observation period, and as a function of stimulus treatment, lizards modified their rates of horizontal looking (ANOVA: *F* = 10.12, *p* < 0.001; Figure 2E), horizontal locomotion (ANOVA: *F* = 6.2226, *p* < 0.001; Figure 2F), vertical looking (ANOVA: *F* = 4.92, *p* = 0.002; Figure 2G), and vertical locomotion (ANOVA: *F* = 6.84, *p* < 0.001; Figure 2H). All stimuli elicited elevated rates of horizontal looking and locomotion relative to control conditions (all *p* < 0.05; Figure 2E). Lizards exhibited increased vertical looking rates when exposed to rabbits (*t* = 2.85, *p* = 0.024; Figure 2G) relative to control conditions (*t* = −3.60, *p* = 0.002; Figure 2G). Similarly, both rabbits (*t* = 4.43, *p* < 0.001; Figure 2H) and cats (*t* = 2.787, *p* = 0.0281; Figure 2H) elicited elevated rates of vertical locomotion compared to snakes, but none differed from the control (all *p* > 0.05; Figure 2H).

## Discussion

As urbanization continues to force wild populations to coexist with synanthropic predators, like cats, it becomes increasingly important to understand the behavioral implications on wildlife and the degree to which native prey are able to assess the risk of such predators. To address this need, we sought to determine whether western fence lizards responded differently to native predators (snakes), synanthropic predators (cats), and native non-predators (rabbits). We found that cats elicited increased horizontal looking behavior in lizards immediately after stimulus exposure, a response not initially elicited by other stimuli, indicating a rapid ability to detect and respond to cats above and beyond other potential threats. By the end of the trial period, all visual stimuli evoked elevated looking behavior relative to control conditions, such that cats were no longer differentiated from other stimuli. Effects of visual stimuli on locomotion were more nuanced. Rabbits and cats elicited higher rates of vertical locomotion behavior than snakes across the complete observation period, but differences from control conditions were not evident early on. Collectively, our findings reveal that lizards distinguish cats from native predators and non-predators immediately following exposure. However, behavioral responses to all visual cues become heightened over time such that cat-specific discrimination is temporary.

Given the substantial contribution of cats to wildlife mortality in urban areas, understanding whether urban prey are capable of responding to cats is important from a conservation perspective. Cats elicited the strongest immediate behavioral response, suggesting that lizards distinguish between cats and other animals upon initial exposure. This evidence is consistent with studies on lizard antipredator responses to cats. In New Zealand, lizards living in areas populated by cats modify their time budgets by reducing basking when exposed to live cats (Cliff et al. 2022). Similarly, lizards in areas of high cat density in Greece remain closer to refuge and exhibit greater flight initiation distance and tail shedding when approached by cat decoys (Li et al. 2014). Together, our findings contribute to growing evidence that threat perception of cats in lizards is widespread, reinforcing the apparent non-lethal effects of cats on urban wildlife behavior.

Because cats have been present in North America for approximately 450 years, they are no longer considered novel predators (Welker et al. 2025). Therefore, urban prey which coexist with cats have been able to develop recognition of cats as a threat (Steindler et al. 2020; Banks et al. 2018; West et al. 2018), particularly in urban environments where cat density is substantially higher (Bogrand et al. 2017; Flockhart et al. 2016). While urban lizards in North America may be impacted by cats (Loss et al., 2013; Lepczyk et. al. 2023), our findings suggest that mortality is not due to their inability to identify cats as predators upon initial exposure. Because urban prey exhibit stronger antipredator responses in areas of high cat density (West et al. 2018), increased exposure may lead to more accurate predator recognition over time. Therefore, high levels of urban wildlife mortality caused by cats may be better explained by cat density rather than predator recognition alone.

The immediate and unique response to cats highlights the importance of studying immediate behavioral changes following exposure to visual cues of predators. Because prey must engage in rapid risk assessment when they visually detect a predator (Cronin 2005), similar work on antipredator vigilance to visual cues has often focused on immediate responses upon detecting the sight of a predator (e.g., Blumstein et al. 2000; Zoratto et al. 2014; Carlson et al. 2017). Consistent with previous findings on antipredator behavior, cat-specific responses were distinct and immediate, but discrimination diminished over time. This pattern may reflect that that visual detection of a predator creates an immediate risk that must be evaluated, ranked according to its threat, and managed.

The shift in behavior elicited by all stimuli demonstrates that animal videos can be effectively used to study how prey may respond to potential predators. Moreover, our study justifies further use of AI tools to advance current approaches of studying antipredator behavior. Historically, taxidermic mounts have been used to study prey responses to predator cues (Tinbergen 1989; Curio et al. 1978). However, recent studies on lizard perception found no significant behavioral responses to taxidermic mounts of cats (Lishmund et al. 2025). In contrast, our dynamic videos of cats triggered behavioral changes not achieved by static mounts, indicating that movement may be crucial for accurate predator recognition. More recently, animated videos and video playback have been used to study antipredator behavior (Gerlai et al. 2009; Watve et al. 2019), which are currently the most realistic substitutions for live animals. Even so, such mechanisms often fail to ensure that behavioral responses do not arise from one singular stimulus depiction. Our creation of multiple exemplars per stimulus type guaranteed that each lizard was exposed to a slightly different depiction of each animal, allowing us to attribute behavioral responses to predator type rather than to a single stimulus depiction.

Future studies of lizard responses to videos of predators, like cats, may benefit from better standardizing the starting position of each individual. We intentionally did not begin trials until the individuals were relaxed and free of disruption, which resulted in variation in lizard starting positions. Some lizards were horizontal (on the substrate), while others were vertical (on the sides of the enclosure) at the time that focal observations began, limiting our ability to identify predator-specific escape behavior. Lizards living with ground dwelling predators have been found to favor higher perches (Pringle et al. 2019), while those living without such predators spent more time in areas lower to the ground, suggesting that presence and perception of predators may influence micro-habitat selection (Lapiedra et al. 2023). By controlling for initial location, future studies could determine whether specific predators drove animals to vertical substrates.

While our study focused on visual predator discrimination, prey rely on multiple modalities when assessing risk (Kats & Dill 2016; Hettena et al. 2014; Caro 2005). Prior studies have found more robust responses when individuals were exposed to multiple modalities, such as observations of heightened responses to visual and chemical cues of snakes relative to a single modality (Jones et al. 2024; Amo et al. 2004). Although lizards perceive dynamic visual representations of cats and adjust behavior accordingly, their inability to discriminate between stimuli over time suggests that visual cues alone may be insufficient when studying cat recognition in urban prey. Integration of additional modalities such as olfactory cues may provide more information on the behavioral effects of cats on lizards as well the ability of urban prey to identify and evade cats.

Overall, our findings indicate that urban fence lizards detect and respond to AI-generated videos of moving cats, but this cat-specific discrimination is short-lived, as responses to visual stimuli of other animals emerge over time. The ability of lizards to rapidly respond to cats indicates that urban prey may not be naive to synanthropic predators associated with urbanization. Ongoing mortality may be more of a consequence of cat density levels or their efficiency in hunting lizards and other small prey. Future work is needed to understand whether cats ‘out-gun’ (Bleicher 2017) urban prey or whether reducing cat densities is essential for coexistence.

## Acknowledgements

We thank Allison Keeney and Prof. Victoria Sork for granting permission to use the UCLA Botanical Garden for our experiments, Dr. Bree Putman for helpful advice on the experimental design, Stephen Hanawalt for assistance with lizard detection and capture, and the Wild Animal Initiative for funding the study.

## Data Availability Statement

Original data and R scripts are archived on Dryad and will be made publicly available at the time of acceptance.

## Funding Statement

The study was funded through a Wild Animal Initiative postdoctoral fellowship held by DRD

## Statement of Author Contributions

EP, DRD, DTB, and HBS collectively developed the idea for this study. EP carried out the experiment, data processing, and data analysis, with support from DTB, DRD, and HBS. EP wrote the initial draft of the manuscript, which was revised and improved with the help of all other co-authors.

## Conflict of Interest

The authors declare no conflict of interest.

## Ethics Approval Statement

All procedures were approved by University of California Los Angeles (UCLA) Institutional Animal Care and Use Committee (Protocol No. ARC-2025-016; held by DRD and HBS) and operated under approved scientific collecting permits California Department of Fish and Wildlife (SC 2480, held by HBS).

---

**Table 1.**
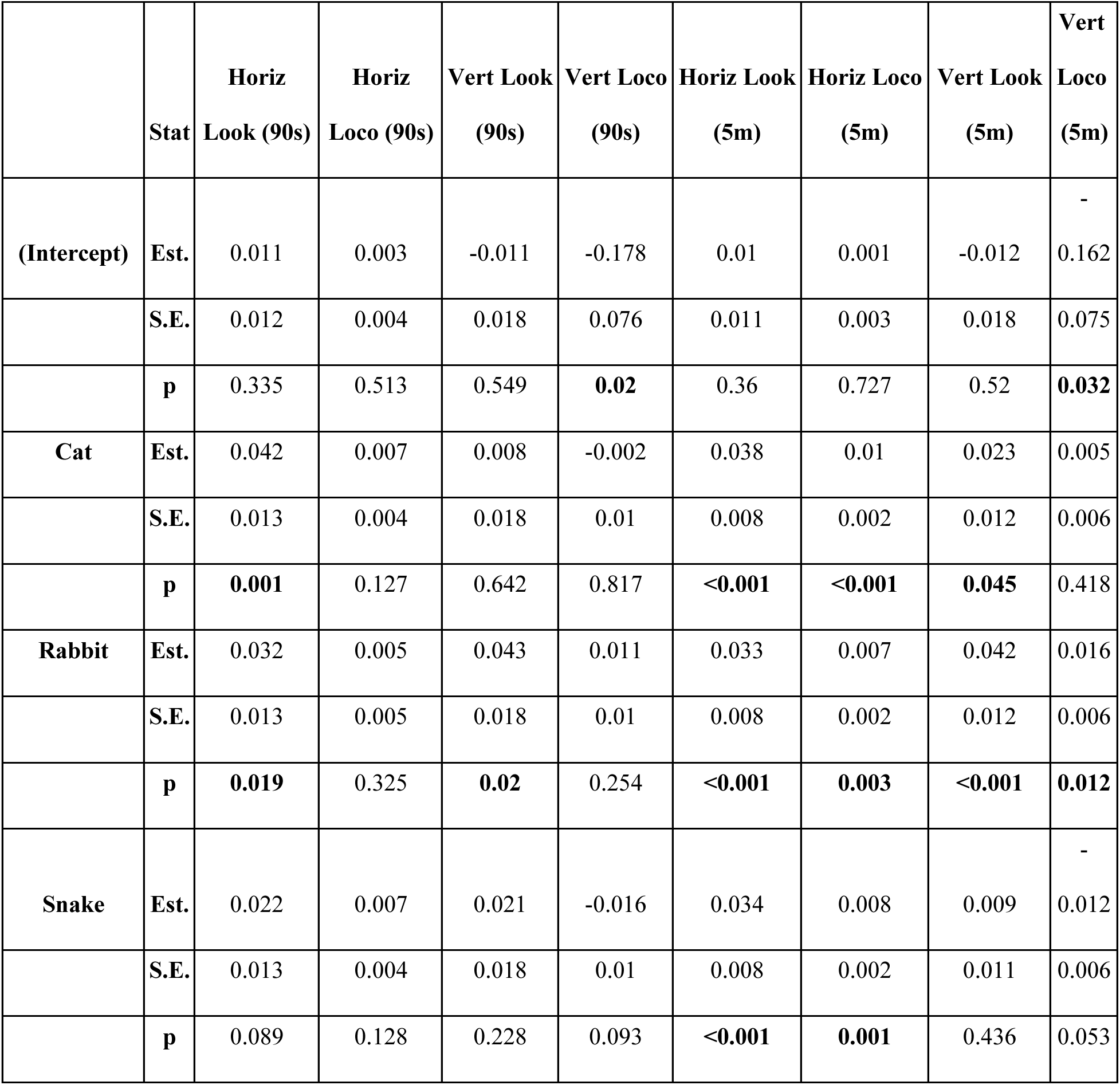
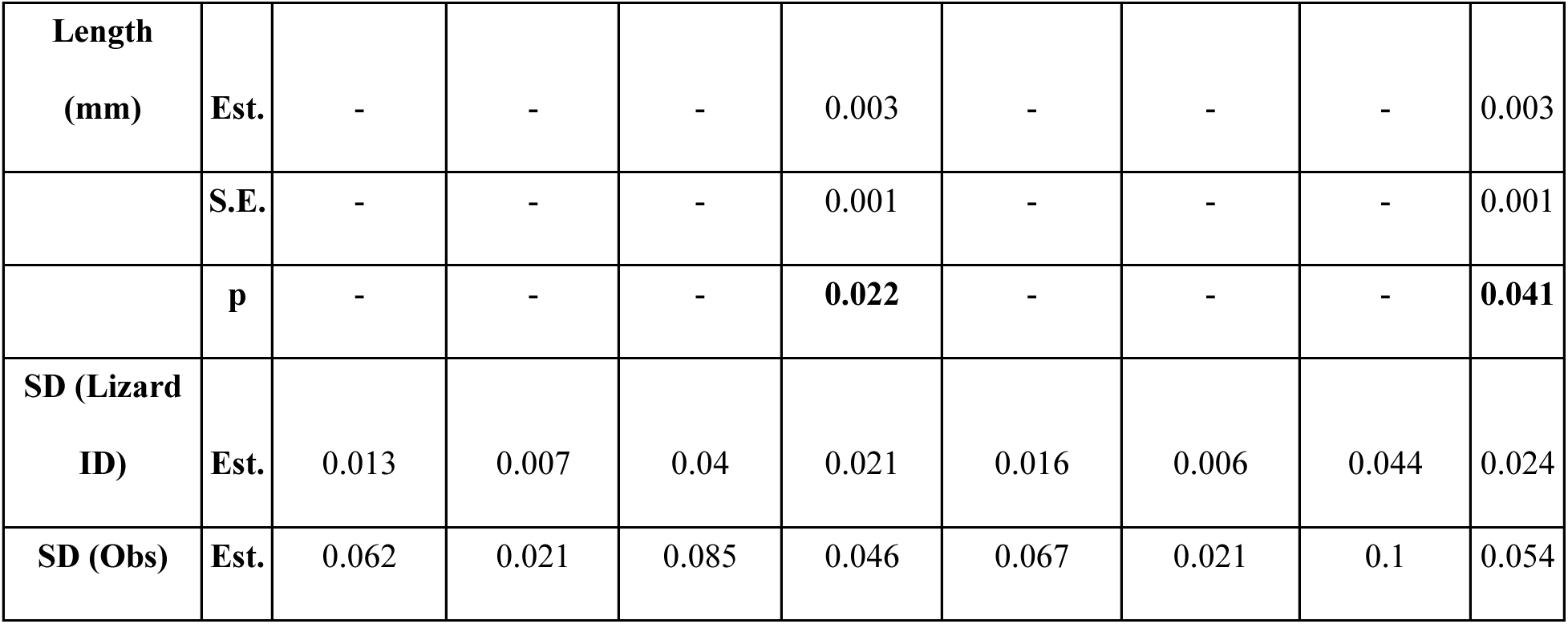
Summary of fixed effects coefficients for models of looking and locomotion behavior during the first 90 s and 5 min post-stimulus.

## Notes

### Competing Interest Statement

The authors have declared no competing interest.

