## Supplementary Material for "Can urban fence lizards identify cats?"

#### Supplementary material for:
### Can urban fence lizards identify cats?

Emma M. Pfister, Daniel T. Blumstein, H. Bradley Shaffer, David R. Daversa

**Table S1.** Latin square trial design implemented for the experiments. The design was replicated four times (N = 16 lizards).

| | **Day 1** | **Day 2** | **Day 3** | **Day 4** |
|---|---|---|---|---|
| **Lizard 1** | Control | Cat | Snake | Rabbit |
| **Lizard 2** | Cat | Snake | Rabbit | Control |
| **Lizard 3** | Snake | Rabbit | Control | Cat |
| **Lizard 4** | Rabbit | Control | Cat | Snake |

**Table S2.** Ethogram used to score lizard behaviors.

| **Behavior** | **Definition** |
|---|---|
| Look | Stationary and quadrupedal, each head turn left/right is an additional look |
| Locomotion | Movement from one location to another |
| Elevated look | Stationary, body positioned closer to the ground, each head turn left/right is an additional look |
| Flattened look | Front legs fully extended, sustained elevation, each head turn left/right is an additional look |
| Push-up | Rapid elevation and lowering of body |

| | |
|---|---|
| Head bob | Movement of head up and down |
| Tongue flick | Rapid outward and inward movement of tongue |
| Out of sight | Fleeing under hiding spot or burrowing (no longer visible) |

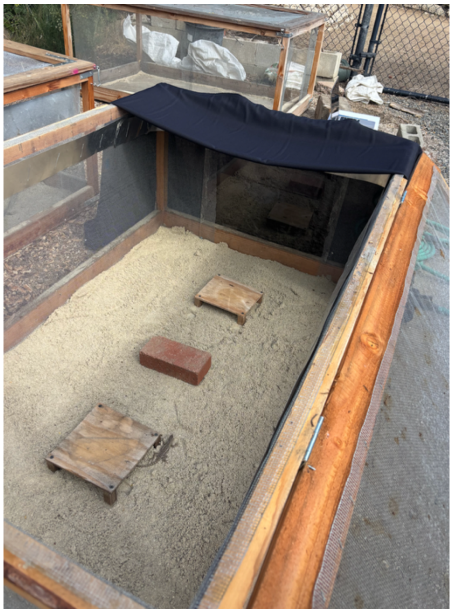

**Figure S1.** Enclosure setup. Wooden frames with the dimensions 5'L x 3'W x 2'H were covered in hardware cloth on all sides. One short side of the cage had a square plexiglass cutout for placing the monitors. Black curtains were placed around the plexiglass side and one long side of the enclosures to reduce glare and prevent lizards from seeing each other. Enclosure enrichment included an ~2 cm layer of sand, two wooden shelters and one brick.

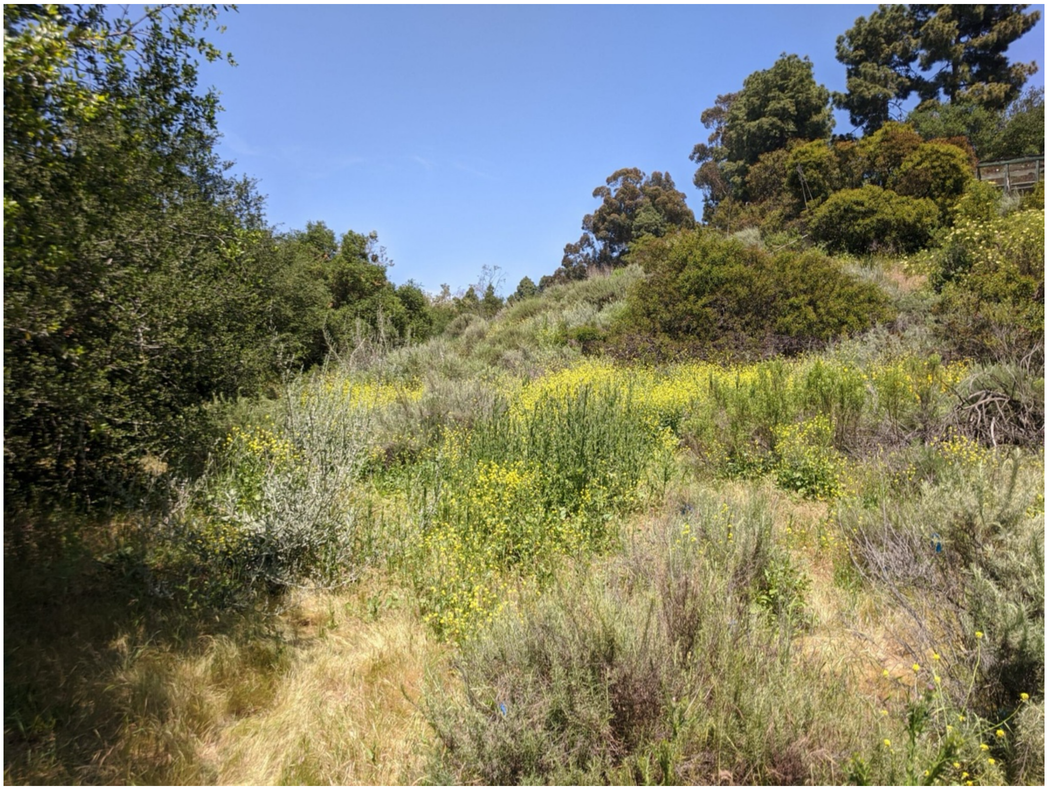

**Figure S2.** Sage Hill background image used for all videos.

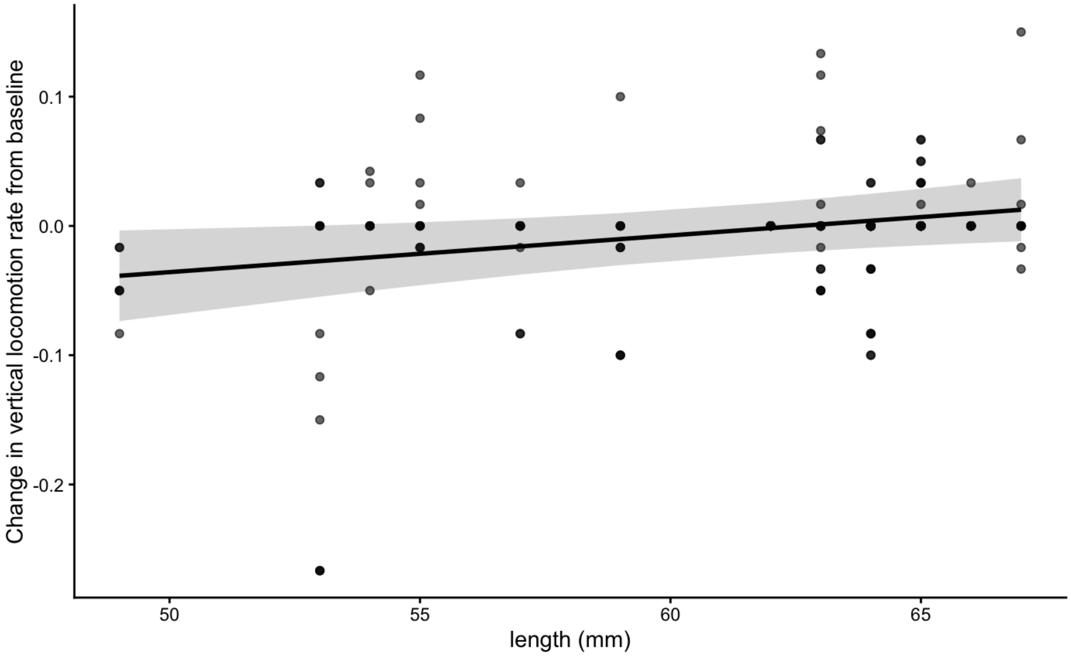


**Figure S3.** Correlation between lizard snout-to-vent length and the change in their vertical look rate from baseline.

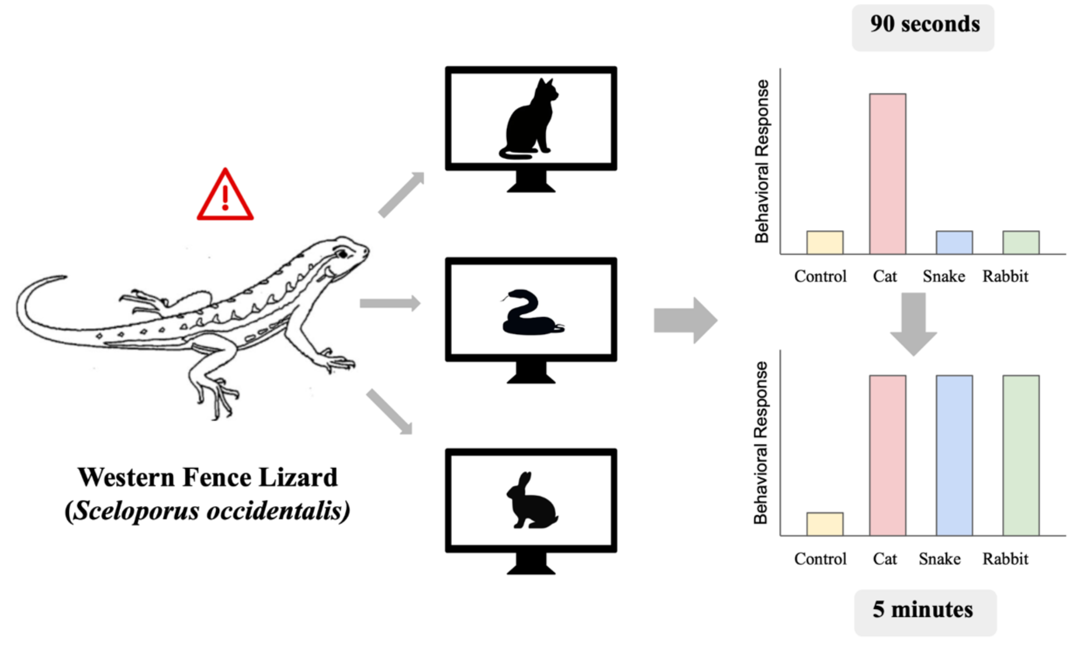


**Figure S4. Graphical Abstract.** Following encounters with artificial intelligence-generated videos of cats, snakes, and rabbits, western fence lizards (*Sceloporus occidentalis*) from the campus of the University of California, Los Angeles, initially discriminated cats from other stimuli in their horizontal looking behavior. The cat-specific looking rates, while still higher than controls, became similar to looking rates at other stimuli by the end of the 5-minute trial. Lizards initially discriminate cats from other animals, but only temporarily.

**Videos S1 - S16.** 15-second stimulus videos used in the experiments (videos attached as separate files).
